# Eukaryotic-driven directed evolution of Cas9 nucleases

**DOI:** 10.1101/2023.09.18.558227

**Authors:** Giulia Vittoria Ruta, Matteo Ciciani, Eyemen Kheir, Michele Domenico Gentile, Simone Amistadi, Antonio Casini, Anna Cereseto

## Abstract

**Full exploitation of the natural reservoir of CRISPR-Cas nucleases from prokaryotes for genome editing is limited by the suboptimal activity of these enzymes in mammalian cells. Here we developed a <u>E</u>ukaryotic <u>P</u>latform to Improve <u>C</u>as <u>A</u>ctivity (EPICA) to steer weakly active Cas9 nucleases into highly active enzymes by directed evolution. The EPICA platform is obtained by coupling Cas nuclease activity with yeast auxotrophic selection followed by mammalian cell selection through a sensitive reporter system. EPICA was validated with a poorly efficient Cas9 nuclease from *Campylobacter jejuni*, CjCas9, generating an enhanced variant, UltraCjCas9, following directed evolution rounds. UltraCjCas9 was up to 12-fold more active in mammalian endogenous genomic loci, while preserving high genome-wide specificity.**

**Here we report a eukaryotic pipeline allowing enhancement of Cas9 systems, setting the ground to unlock the multitude of RNA-guided nucleases existing in nature.**

## INTRODUCTION

The development of CRISPR-Cas tools (1, 2) and the emerging number of RNA-guided prokaryotic and eukaryotic nucleases (3–5) are giving a tremendous impulse to the development of advanced therapies based on genomic modifications (6). Yet, to further accelerate and broaden the application of genome editing (7), a larger set of tools are highly needed to match the complexity of gene therapy approaches. In particular, there is a demand to increase the accessibility to any genomic site by enlarging the repertoire of enzymes covering the largest number of protospacer adjacent motifs (PAM) (8, 9). Moreover, the efficacy of cellular and organ delivery, which highly depends on the cargo size (10), is very relevant to the field, thus reinforcing the need for low molecular weight Cas enzymes. Small molecular size is indeed necessary for Cas compatibility with adeno-associated viral vector (AAV) without enzyme splitting (11) and allows the generation of compact base editors and prime editors, which can precisely edit DNA avoiding double-strand breaks (1, 2).

A large reservoir of CRISPR-Cas systems and novel emerging ancestral RNA-guided nucleases have been identified in metagenomic data banks. Nonetheless, the majority of enzymes isolated from bacteria (12), archaea (13) and viruses (14) are poorly active in mammalian cells. Strategies aimed at enhancing these systems for genome editing include molecular engineering of the nucleases by rational design (15), bacterial screening (16) or optimization of the gRNA (17). Despite these efforts, few CRISPR-Cas tools are sufficiently active for efficient genome editing in eukaryotes (18), as demonstrated by the paucity of RNA-guided nucleases for experimental use, as opposed to the multitude of systems identified in metagenomic data banks. To unlock the natural reservoir of prokaryotic systems, experimental setups are needed to enable the adaptation of enzymes to the eukaryotic nuclear environment. We have previously demonstrated that yeast cells are ductile eukaryotic systems that can improve Cas precision by directed evolution (19). On this groundwork we generated a eukaryotic platform to improve Cas activity (EPICA). To demonstrate the efficacy of this evolution platform, we introduced in EPICA CjCas9 (20), a small size (984 aa) Cas enzyme with favorable features for genome editing, which is however minimally used due to its reduced nuclease activity (21). The platform produced an enhanced variant, UltraCjCas9, carrying modifications hardly predictable through a rational engineering approach.

## RESULTS

### Yeast based directed evolution enhances Cas9 activity

To generate a eukaryotic platform for the evolution of Cas9 nuclease activity, a yeast strain was engineered by integrating two different cleavage reporter cassettes in the tryptophan (TRP) and adenine (ADE) genes. Disruption of the TRP and ADE genes by cassette insertion turns yeast growth dependent from Adenine and Tryptophan culture integration. The cassettes contain target sites including different PAM sequences compatible with the Cas9 under evolution, to allow the isolation of variants evolved independently from specific PAM sequence recognition. Homology arms were inserted at the edges of the cassettes to mediate single strand annealing (SSA) repair induced by Cas9 cleavage of the target locus leading to gene reconstitution (19). This model, schematized in **Figure 1a**, allows selection of active Cas9 molecules by auxotrophy through limiting growth factors conditions (TRP and ADE deprivation, TRP- and ADE- respectively), permitting exclusive proliferation of cells with restored TRP and ADE loci generated by efficient Cas9 cleavage. The platform was initially tested with SpCas9, showing high levels of yeast cell survival in single ADE- and TRP- or combined ADE-/TRP-; by contrast, wild-type CjCas9 (CjCas9 WT) showed 20% of living colonies in ADE- conditions but nearly absent survival in ADE/TRP deprived culture conditions (**Fig 1b**). Transformation of a cDNA library of CjCas9 variants obtained through random PCR mutagenesis led to isolation of colonies growing in ADE-/TRP- conditions (**Fig. 1c**). Starting from the colonies obtained by transformation of the mutagenized CjCas9 library, the process of random mutagenesis and transformation in ADE-/TRP- conditions was reiterated until a decreasing trend in cell survival was observed (rounds 5 and 6), likely resulting from saturating mutagenesis. The cDNA isolated from the fourth round of mutagenesis, where the highest degree of cleavage activity was detected (227-fold compared to wild-type, **Fig 1c**), was deep sequenced revealing a large variety of substitutions throughout the nuclease sequence (**Fig. 1d**). As an intermediate step to verify the yeast evolution step, we sampled the cDNA library to evaluate genome editing improvement in mammalian cells. We selected the most frequent substitutions (E189G, F214I, E789K, T913S, **Fig. 1d**) which were tested individually for indel formation in three endogenous loci (*RHO*, *CCR5* and *EMX*) of HEK293 cells. Each single mutation showed variable editing improvement (**Fig. 1e**), thus suggesting that enhancement in yeast corresponded to optimization in a mammalian context.

**Figure. 1.**
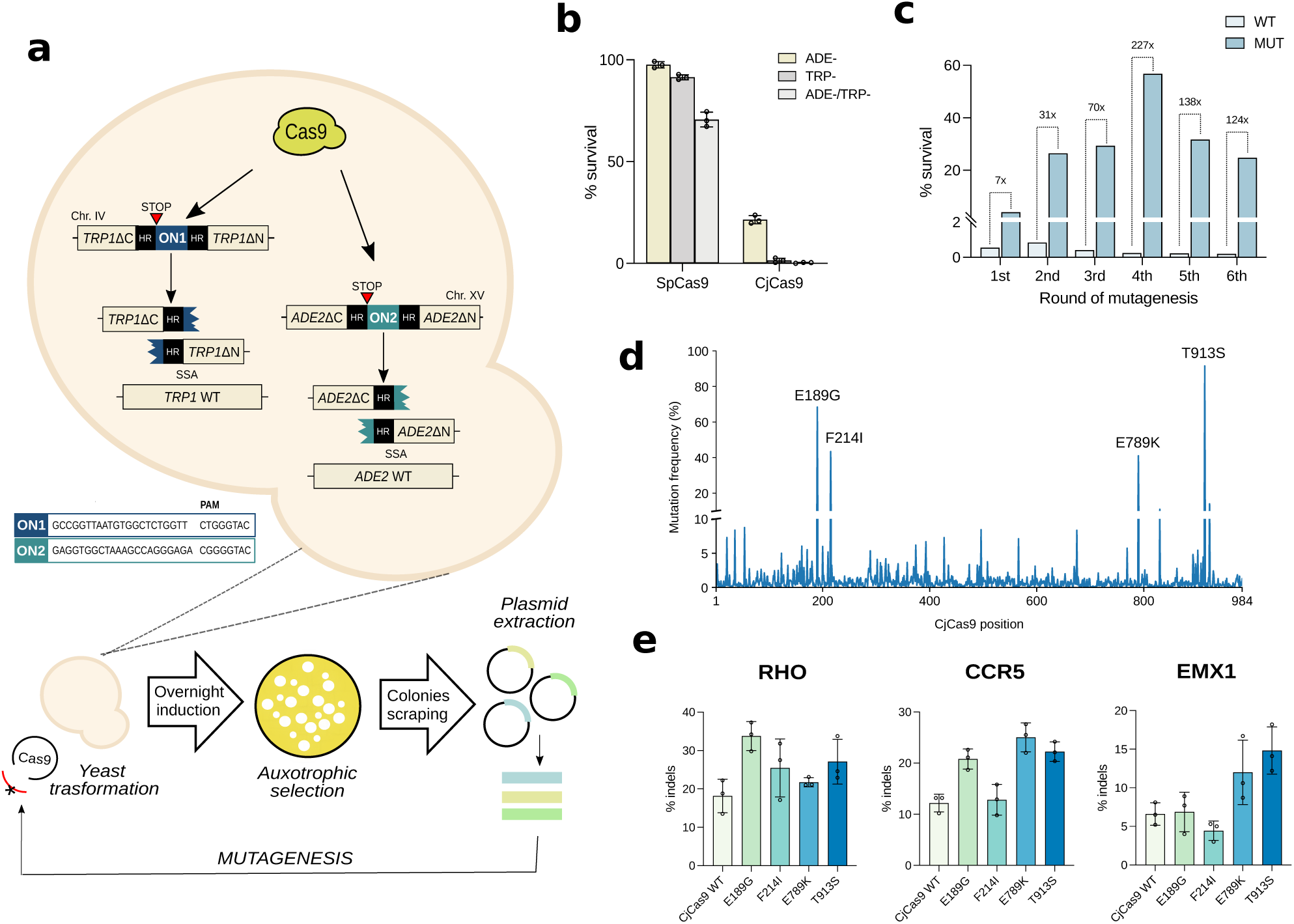
Directed evolution step to enhance Cas nuclease activity. (**a**) Schematic and experimental flowchart of the yeast Cas evolution step. Framed sequences correspond to the ON1 and ON2 cassettes carrying the CjCas9 target sites and AAVS1-TS34 and AAVS1-TS32 targets, respectively. Active mutants cleaving both ON1 and ON2 in the ADE2 and TRP1 loci are amplified from colonies growing in ADE- and TRP- conditions and undergo additional rounds of mutagenesis and selection. (**b**) Cleavage activity of SpCas9 and CjCas9 are reported as percentages of survival, corresponding to the number of colonies growing in TRP- and ADE- plates with respect to total transformants measured in unrestricted growth conditions. Culture plates shown in **Supplementary Figure 1a**. (**c**) The activity of the CjCas9 variants (MUT) isolated at each round of evolution were compared with CjCas9 WT by measuring the percentages of survival after transformation. Culture plates shown in **Supplementary Figure 1b**. (**d**) Peaks at each CjCas9 amino acid position represent the frequency of the mutated residues obtained by NGS deep sequencing. Amino acid substitutions with high frequency are reported in the graph. (**e**) Editing activity at endogenous loci in HEK293 cells of CjCas9 variants containing substitutions corresponding to the most frequently mutated amino acids in (**d**). In (**b**) and (**e**) data are reported as mean standard deviation of n≥3 biologically independent samples. Individual values are represented as empty circles.

### Selection of the enhanced CjCas9 variants in a mammalian reporter cell line

To select the best combinations of mutations for CjCas9 enhancement in mammalian cells, we generated a reporter cell line carrying a switch-on EGFP circuit controlled by indel formation. HEK293 cells carrying a Tet repressor (TetR) cDNA were engineered with a EGFP cassette repressed by Tet operator (TetO) elements, which are released by CjCas9 cleavage of TetR (**Fig. 2a** and **Supplementary Fig 2a**). The Cas mediated induction of fluorescence increases sensitivity compared to the widely used EGFP disruption assay (19), thus facilitating detection of differences in Cas activity (**Supplementary Fig 2a)**. A lentiviral library of the yeast-evolved CjCas9 variants was transduced in the TetR-EGFP reporter cell line to select the most active mutants (**Supplementary Fig. 2b**). The selection step was repeated a second time to further enrich the best variants, which showed almost 10-fold higher fluorescence than controls, indicating the presence of mutants more active than the original enzyme (**Fig. 2b** and **Supplementary Fig. 2b**). A large number of variants (n=160) were isolated from double-sorted EGFP cells and tested individually (**Supplementary Fig. 2c**). We selected the best performing variants, corresponding to mutants having the highest ratio of positive EGFP cells with respect to CjCas9 WT in the TetR-EGFP reporter cell line. The best performing variants (n=17) were further evaluated for indel formation in the *EMX1*, *RHO* and *CCR5* endogenous loci (**Fig. 2c**), revealing an average increased activity compared to wild-type ranging from 2 up to 3.6-fold (**Supplementary Table 1**).

**Figure. 2.**
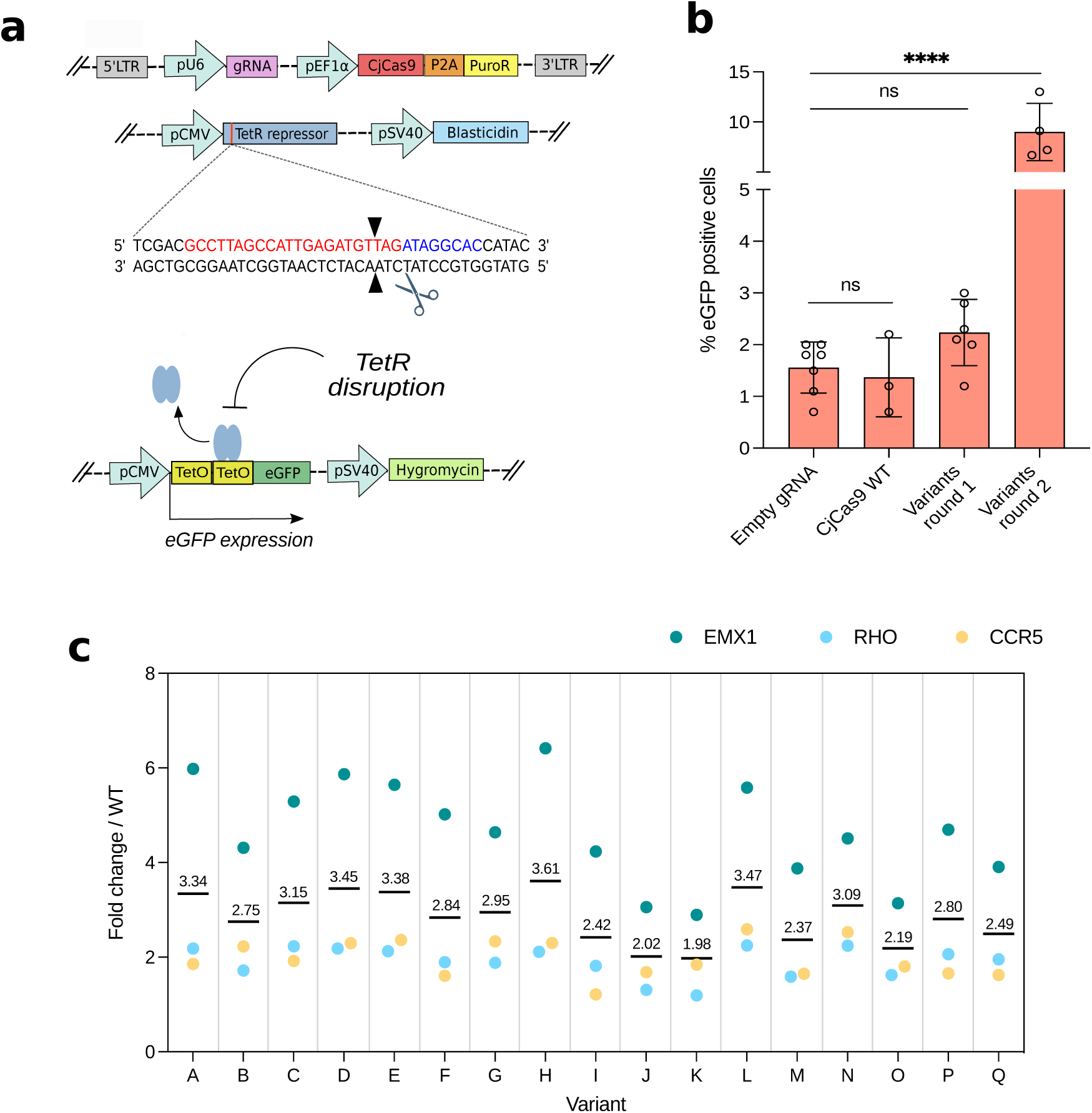
CjCas9 enhancement through screening in mammalian cells. (**a**) Schematic representation of the TetR-EGFP reporter cell line. TetR (blue) is constitutively expressed inactivating EGFP (green) expression. TetR inactivation by Cas9 cleavage enables EGFP expression and isolation of fluorescent cells by sorting. (**b**) CjCas9 variants isolated from rounds 1 and 2 of the mammalian screening are compared with CjCas9 WT transducing the TetR-EGFP reporter cell line. Statistical analysis was performed using one-way ANOVA with Tukey’s multiple comparisons test: ns not significant, **** P < 0.0001. (**c**) Editing activity of selected variants obtained from yeast and mammalian selection. Dots represent fold changes of activity of each variant (from A to Q) with respect to CjCas9 WT targeting EMX, RHO or CCR5. Extended data shown in **Supplementary Figure 3**. Grand mean values of each variant are shown in the graph. Sanger sequences of the variants are reported in **Supplementary Table 2**. In (**b**) data are reported as mean ± standard deviation of n≥3 biologically independent samples. Individual values are represented as empty circles.

In conclusion, through the evolution pressure obtained by coupling Cas cleavage activity with auxotrophic selection in yeast and indel formation in mammalian cells, EPICA enables the generation of enhanced Cas variants.

### UltraCjCas9, an evolved highly active Cas9

The most active variant, named UltraCjCas9 hereafter, was further characterized. Sanger sequencing revealed five modifications in four domains (RuvC-I, REC1, HNH and PI) (**Fig. 3a**). To evaluate the editing profile we performed a comparative editing analysis in 23 human genomic loci with UltraCjCas9, CjCas9 WT and the recently optimized enCjCas9 (15) showing that UltraCjCas9 was more active in the majority of sites (**Fig. 3b**). The editing efficacy of UltraCjCas9 was quite variable, reaching up to 12-fold higher activity than the wild-type (*HEK site 1*) (**Fig. 3b** and **Supplementary Table 3**). Overall UltraCjCas9 was more active than CjCas9 WT and outperformed enCas9, which was obtained by rational engineering (**Fig. 3c**). Given the editing efficacy of UltraCjCas9, we compared this variant with the most commonly used Cas9 from *Streptococcus pyogenes*, SpCas9, by selecting genomic loci efficiently edited by UltraCjCas9 and using overlapping guides for both orthologs (**Supplementary Fig. 5b**). Overall, the two nucleases showed no significant difference in editing efficiency (**Fig. 3d**).

**Figure. 3.**
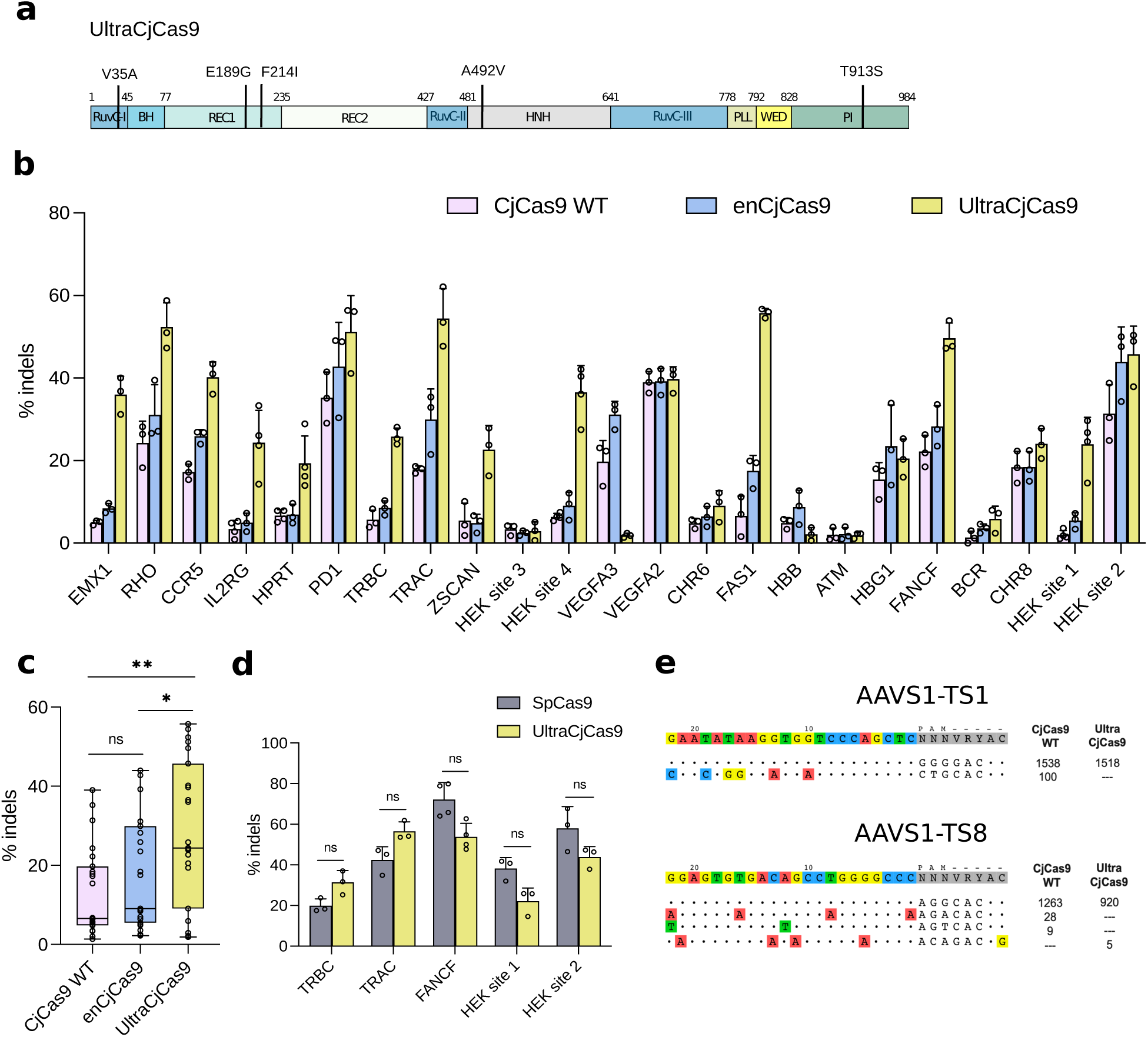
Characterization of the highly enhanced variant UltraCjCas9. (**a**) Domain localization of the five amino acid substitutions (V35A, E189G, F214I, A492V and T913S) characterizing UltraCjCas9. See **Supplementary Figure 4** for amino acid positions in the CjCas9 structure. (**b**) Editing efficiency in endogenous genomic loci of HEK293 cells with CjCas9 WT, enCjCas9 and UltraCjCas9. PAM sequences reported in **Supplementary Figure 5a**. (**c**) Graphical summary of editing activities of CjCas9 WT (pink), enCjCas9 (blue) and UltraCjCas9 (yellow) in (**b**). Empty circles in the box plots represent the average percentages of indels for each genomic locus. Statistical analysis was performed using one-way ANOVA with Tukey’s multiple comparisons test: ns not significant, * P < 0.05, ** P <0.01. Central line, median; box limits, upper and lower quartiles; whiskers, ×1.5 interquartile range; n=23 independent loci. (**d**) Comparison of SpCas9 and UltraCjCas9 editing activity in the indicated endogenous genomic loci. Statistical significance was assessed using a two-sided t-test corrected using the Holm-Šídák method for each locus. ns not significant. (**e**) GUIDE-Seq analysis performed with CjCas9 WT and UltraCjCas9. GUIDE-seq read counts of each on- and off-target in HEK293T cells are shown on the right side (Extended data in **Supplementary Tables 7-10)**. In (**b**) and (**d**) data are reported as mean ± standard deviation of n≥3 biologically independent samples. Individual values are represented as empty circles.

Finally, since enhanced Cas9 nuclease activity may correlate with decreased fidelity (22), we performed a whole-genome off-target analysis by GUIDE-Seq (23) to compare the specificity of CjCas9 WT and UltraCjCas9. We chose two sites, *AAVS1-TS1* and *AAVS1-TS8*, previously associated with CjCas9 off-target activity (20), where both nucleases produced similar levels of on-target indels (**Supplementary Fig. 5c**). In both genomic loci UltraCjCas9 generated less off-target cleavage sites than the wild-type enzyme, resulting in slightly higher precision (**Fig. 3e**). Overall UltraCjCas9, generated by EPICA, showed an enhanced activity while preserving specificity.

## DISCUSSION

In just over a decade, CRISPR-Cas systems working as adaptive immunity in prokaryotes have been successfully translated into genome editing tools for mammalian cells up to the clinic. Notably, the prokaryotic cellular environment highly differs from the eukaryotic nuclear compartment, characterized by multiple factors defining chromatin structure and function. Hence, to further advance this technology, CRISPR-Cas systems should be functionally steered in eukaryotic cells. Along this line, emerging data from various groups demonstrated that chromatin structure impacts CRISPR-Cas genome editing efficacy (24–32). Remarkably, while nucleosomes affect the editing activity of CRISPR-Cas enzymes, the artificial zinc finger nucleases (ZFN) remain unaffected, suggesting a natural evolution of Cas nucleases to work with naked DNA (27). More recently, direct evidence demonstrated the role of chromatin structure on Cas12a PAM recognition and R-loop formation efficiency (30) and the effect of chromatin remodeling factors on the nuclease activity of a variety of Cas9 proteins, including small orthologs (31). While the effect of chromatin structure on Cas efficiency has been well documented, other factors could influence nuclease activity in eukaryotic cells, such as interactions with host proteins. Post-translational modifications (PTMs) have been shown to inhibit Cas proteins in prokaryotes (32–34), however it remains unclear to what extent nuclease activity could be modulated by PTMs introduced by host factors in eukaryotes (35). These results suggest that limited insights on Cas interactions with the eukaryotic environment will severely limit the molecular engineering of these tools. While rational design has been widely employed to engineer Cas nucleases (15, 36–38), this approach depends on protein structure information which is not necessarily available for the large number of RNA-guided nucleases requiring optimization as genome editing tools. Moreover, resolved protein structures are often missing flexible regions, such as the HNH domain of Cas9 nucleases (39–41), imposing limitations on the rational design approach. The advantage of an unbiased *in cellulo* Cas9 selection is supported by our results showing the superiority of UltraCjCas9 over the rationally engineered enCjCas9 (15). Remarkably, the mutations in UltraCjCas9 obtained via EPICA would have been hardly predictable through structure analysis. Indeed, mutations V35A, A492V and T913S are unexpectedly chemically similar to the wild-type residues, and mutation F214I is positioned far from DNA-RNA interactions (**Supplementary Fig. 4**). Conversely, enhancement of Cas9 nucleases by rational engineering is generally obtained through modifications of residues directly contacting the DNA-RNA hybrid, often increasing positive charges in order to facilitate interactions between the Cas9 protein and the negatively charged nucleic acid backbone (15, 37). In line with this type of enhancing modifications, UltraCjCas9 carries the substitution E189G, which neutralizes a negative charge in the vicinity of the PAM-proximal region of the DNA-RNA hybrid. Interestingly, modifications introduced in UltraCjCas9 did not interfere with the nuclease specificity, as opposed to previous molecular engineering approaches (37), implying that enhanced nuclease activity does not necessarily increase tolerance to sgRNA-target DNA mismatches for Cas9 proteins, as also previously observed for CasΦ (42).

Here we report a fully eukaryotic platform for Cas enhancement. Our methodology exploits evolutionary steps in yeast followed by selection of variants in mammalian cells. Enhanced nucleases such as UltraCjCas9, obtained through EPICA, will foster the expansion of the genome editing toolbox with the development of a larger variety of CRISPR-Cas systems. Since the platform allows the evolution of enzymes with nuclease activity, its use can be potentially extended beyond CRISPR-Cas systems, including the novel emerging collection of RNA-guided nucleases associated with prokaryotic and eukaryotic transposons (3–5, 43). The modification of Cas9 proteins steered under the pressure imposed by the complexity of the nuclear environment may shed light on critical residues required for nuclease enhancement in eukaryotes, thus facilitating the translation of prokaryotic resources for genome editing applications.

## MATERIALS AND METHODS

### Plasmids

The constructs used for the generation of the yeast strain were obtained by cloning the AAVS1-TS34 and AAVS1-TS32 targets and PAM sequences as annealed oligonucleotides in pUC19 plasmid containing respectively ADE and TRP cassettes, after digestion with KpnI and BamHI. For expression in yeast, the sequence of CjCas9 was amplified from the pX404 (Addgene 68338) and cloned in place of SpCas9 in the LEU2 carrying plasmid p415-GalL-Cas9-CYC1t (Addgene 43804), by double digestion with SpeI/XhoI. pSNR52-BsmBI-Cj_sgRNA for gRNA expression in yeast was generated by substituting the promoter in the pU6 Cj sgRNA plasmid (Addgene 89753) with the SNR promoter by double digestion with XhoI and BamHI. The AAVS1-TS34 and AAVS1-TS32 target sequences were cloned as annealed oligonucleotides in the pSNR52 Cj gRNA digested with BsmBI. The SNR- AAVS1- TS34 and -TS32 Cj gRNA fragments were first cloned in the URA3 carrying pRS316 plasmid by double digestion with XhoI/SacII adding the SUP4t yeast terminator; for double guide plasmid the SNR- AAVS1TS32 Cj gRNA fragment was then amplified and cloned in the generated pRS316-SNR52p-AAVS1TS34gRNA -SUP4t plasmid by digestion with SacI. The Sp sgRNA plasmids were obtained through PCR- site directed mutagenesis of p426- SNR52p-gRNA.CAN1.Y-SUP4t (Addgene 43803) to introduce the target sequences. For mammalian expression the CjCas9 gRNA scaffold was optimized through site-directed mutagenesis to interrupt the A/T stretches (31); targets were cloned as annealed oligos in the pU6 opt CjCas9 gRNA cut with BsmBI. List of gRNAs cloning primers is available in **Supplementary Table 4**. For expression in mammalian cells, CjCas9 WT was cloned in a pX330 containing the chicken β-actin promoter by double digestion with AgeI and EcoRI/BbsI. To generate single mutants, CjCas9 wildtype sequence was cloned in a pEGFP-N1 with AgeI and MfeI and single substitutions introduced through PCR-site-directed mutagenesis; the protein sequence was then cloned back in pX330. Multiple mutants were amplified from the two-rounds enriched sorted cells and cloned in pX330 downstream of a SV40 NLS sequence previously inserted in the plasmid, as well as CjCas9 WT, digesting with KpnI and EcoRI/BbsI. enCjCas9 was generated as double mutant introducing L58Y and D900K mutations in the CjCas9 WT sequence through site directed mutagenesis and then cloned in the pX330 NLS plasmid. Primers used for plasmid cloning are reported in **Supplementary Table 5.**

### Yeast culture

Yeast strains were grown in YPDA-rich medium; for auxotrophic selection yeasts were grown in synthetic minimal medium (SD), omitting the single amino acids required for the experimental purpose. Yeasts stably expressing CjCas9 gRNA were kept growing in SD without uracil before transformation. Cas9 expression was obtained using 20 g/L D-(+)- galactose and 10 g/L D-(+)-raffinose instead of dextrose.

### Cell culture

HEK293 and HEK293T from the American Type Culture Collection (ATCC) were cultured in Dulbecco′s modified Eagle′s medium (DMEM; Life Technologies) with 10% FCS (Life Technologies) and antibiotics (Life Technologies). The EGFP T-REx 293 cell line used for the mammalian screening was obtained modifying the Flp-In™ T-REx™ 293 Cell Line (Invitrogen) for the expression of inducible EGFP according to the manufacturer’s protocol. The cell lines were verified for the absence of mycoplasma contamination (PlasmoTest, Invitroogen).

### Yeast screening

The yeast strain used for the screening was generated from the yLFM-ICORE strain by Delitto Perfetto approach (44), using the constructs pUC19-Ade2-AAVS1TS34 and pUC19-Trp1-AAVS1TS32. The library of CjCas9 variants was produced by random mutagenesis of the wild-type sequence using error-prone PCR approach (GeneMorph II kit from Agilent) (for primers, see **Supplementary Table 5**). The manufacturer’s conditions were followed to set a rate of 0–4.5 mutations/kb; yeasts were co-transformed with the PCR library and the previously digested (SpeI/XhoI) p415-GalL-CYC1t plasmid in a 3:1 ratio, to obtain direct assembly *in vivo*. Yeasts were transformed following the Gietz *et al*. published protocol (45). After transformation, yeasts were grown for 5 hours in SD medium lacking uracil and leucine for recovery and recombination, then the expression of Cas9 was induced through overnight incubation in galactose-containing medium. The next day, yeasts were plated in medium lacking adenine and tryptophan, to select the colonies with enhanced variants. Colonies were collected and the plasmids were extracted with Zymoprep Yeast Plasmid Miniprep II (Zymo Research). For the next rounds of mutagenesis, high fidelity PCR was performed on the selected variants with Phusion™ High-Fidelity DNA Polymerase (Thermo Scientific) and used as template for the subsequent step of mutagenesis (described above). For each round of mutagenesis transformation and collection of selected colonies were repeated. For the validation of the yeast strain, cells were transformed with the entire p415 plasmid expressing wild-type SpCas9 or CjCas9, using the same conditions of the screening. Yeast colonies were counted using the ImageJ software, manually setting the threshold for the maximum discrimination of the colonies. Each yeast experiment was performed by plating a defined amount of yeast suspension in n=3 selective plates as technical replicates.

### Deep sequencing

Variants of the fourth round of mutagenesis were PCR amplified with Phusion High-Fidelity DNA Polymerase (for primers, see **Supplementary Table 5**); the amplicon was used to generate a library with Nextera DNA Flex Library Preparation Kit (Illumina) and sequenced in a PE150 run using Illumina Miseq Reagent kit V2. For data analysis, reads were trimmed using TrimGalore (46) version 0.6.6 (with parameters -q 30 –paired) and aligned to the CjCas9 coding sequence using bowtie2 (47) version 2.3.4.3. Pileups were generated using PaCBAM (48) version 1.6.0 and a custom python script was used to analyze mutation frequencies.

### Indels analysis at human genomic loci

For the evaluation of genome editing activity, 10^5^ HEK293 cells were seeded in 24-wells plates and transfected the next day with 200 ng of gRNA plasmid and 400 ng of Cas9 plasmid; TransIT-LT1 (Mirus Bio) was used as transfection reagent according to manufacturer’s protocol. Cells were collected after 3 days and lysed with QuickExtract DNA Extraction Solution (Lucigen): endogenous loci were PCR amplified with HOT FIREPol MultiPlex Mix (Solis BioDyne), Sanger sequenced and analyzed with the TIDE tool (49). List of TIDE analysis primers are available in **Supplementary Table 6**.

### Lentiviral vector library construction

The CjCas9 gRNA targeting the TetR repressor was amplified from the pU6 plasmid and cloned in the LentiCRISPR V1 by double digestion with NdeI and EcoRI. LentiCRISPR V1 gRNA TetR was then modified to have an intron upstream of the cloning site of Cas9, to avoid a leaky expression of the CjCas9 variants in bacteria and the potential loss of clones. An intron previously used for the same purpose (50) was cloned by insertional mutagenesis through PCR in the middle of the CjCas9 NLS sequence, between a splicing donor site and a splicing acceptor site: the construct Kozak-NLS-V5-intron was then amplified adding a KpnI site downstream and was cloned in LentiCRISPR using the restriction enzymes BamHI and NheI. To minimize the number of background colonies with undigested plasmid, the bacterial toxin ccdB was cloned with the enzymes KpnI and NheI using a dedicated commercial ccdB-resistant strain (ThermoFisher); the generated plasmid LentiCRISPRV1-gRNA-TetR-intron-ccdB was used to obtain the KpnI/NheI digested backbone for lentiviral library. CjCas9 variants were amplified from the plasmid extract of the fourth round of the yeast screening, digested with KpnI and NheI and ligated at room temperature for 2 hours with the digested backbone. The ligation was purified by NucleoSpin Gel and PCR Clean-up (Macherey-Nagel) and used for the electroporation of ElectroMAX DH5α-E Competent Cells (Invitrogen): bacterial colonies were collected until a coverage of at least 100x was reached. The number of different variants was estimated to be 1398, based on the total number of colonies collected from the fourth round of the yeast screening. Pooled library plasmids were purified using NucleoBond PC 500 Maxi kit (Macherey-Nagel). The same protocol was applied to generate the library for the second round of selection, amplifying the variants from the genomic DNA of first-round sorted cells. Primers for the amplification were lengthened for each library to avoid the carry-over of unspecific products from the previous cloning. Primers used for lentiviral vector cloning are reported in **Supplementary Table 5.**

### Cell transduction and sorting

For the lentiviral library production, 2 x 10^7^ HEK293T cells (American Type Culture Collection) were transfected with 25 μg of LentiCRISPR gRNA TetR intron CjCas9 library, 16,2 μg of pCMV-deltaR8.91 and 8,7 μg of VSV-G using the polyethylenimine (PEI) method. Lentiviral particles were filtered through a 0.45 μm PES filter and concentrated with ultracentrifugation. Vectors were resuspended in Opti-MEM (Gibco) and conserved at -80 °C. MOI was estimated by transducing a defined cell number with growing volumes of lentiviral particles and evaluating their viability in puromycin (1 μg/m) after 4 days using an MTT assay. For the rounds of sorting, EGFP T-REx 293 cells were seeded and the next day were transduced with the vector library at MOI∼0.3, ensuring a >100x coverage of the library variants. Cells were then selected in puromycin (1μg/ml) 2 days after transduction and sorted for their EGFP fluorescence after 6 additional days using FACS ARIA III (BD Biosciences). Cells were grown for recovery and collected for genomic DNA extraction with DNeasy Blood and Tissue kit (Qiagen).

### GUIDE-Seq

GUIDE-Seq was performed following a previously published protocol with minor modifications (23). HEK293T cells (ATCC) were transfected with 250 ng of gRNA plasmid, 500 ng of Cas9 plasmid and 10 pmol of dsODNs, using Lipofectamine 3000 (Invitrogen). Cells were selected in puromycin (1 μg/ml) the next day and after 4 days genomic DNA was extracted using DNeasy Blood and Tissue kit (Qiagen). Genomic DNA was sonicated to an average size of 500 bp using S2 Focused-ultrasonicator (Covaris) and end-repaired with NEBNext Ultra End Repair/dA Tailing Module. Adaptors were ligated using NEBNext Ultra Ligation Module as previously described (51). After PCR amplifications, libraries were quantified by Qubit dsDNA High Sensitivity Assay kit (Invitrogen) and sequenced with the MiSeq sequencing system (Illumina) using Miseq Reagent kit V2-150PE.

### CjCas9 structure

The crystal structure of CjCas9 WT in complex with its sgRNA and target DNA was retrieved from PDB (accession 5X2H) (39). To investigate mutations in missing regions, the AlphaFold (52) predicted structure was retrieved from the AlphaFold Protein Structure Database (accession Q0P897) (53) and was aligned to the crystal structure using the Dali server (54). Protein structures were visualized using ChimeraX (55).

### Statistical analysis

Statistical significance tests were performed using GraphPad Prism (version 9.4.1). Data in **Figure 1e** and **Figure 2c** were analyzed with a one-way ANOVA followed by Tukey’s multiple comparisons test. Data in **Figure 2d** were analyzed using two-sided t-tests corrected for multiple comparisons using the Holm-Šídák method. For all analyses, adjusted p-values less than 0.05 were considered statistically significant.

### Data, Materials, and Software availability

The data that support the findings of this study are available from the corresponding author upon reasonable request.

## ACKNOWLEDGEMENTS

We are grateful to Cereseto’s lab for helpful discussion throughout the project. We thank the Next Generation Sequencing facility at the University of Trento for technical support.

This work was supported by the European Union’s Horizon 2020 innovation programme through the UPGRADE (Unlocking Precision Gene Therapy) project (grant agreement No 825825) and Horizon Europe EIC Pathfinder programme AAVolution (grant agreement 01071041).

## AUTHOR CONTRIBUTIONS

G.V.R., E.K., M.D.G. and S.A. designed and performed the experiments; M.C. performed the computational analyses and analyzed the data; G.V.R, M.C., A.Ca. and A.Ce. wrote and edited the paper; A.Ce., A.Ca., conceived and designed the study; A.Ce. was responsible for the coordination of the study. All authors read, corrected, and approved the final manuscript.

## REFERENCES

1. N. M. Gaudelli, et al., Programmable base editing of A•T to G•C in genomic DNA without DNA cleavage. Nature 551, 464–471 (2017).

2. A. V. Anzalone, et al., Search-and-replace genome editing without double-strand breaks or donor DNA. Nature 576, 149–157 (2019).

3. H. Altae-Tran, et al., The widespread IS200/IS605 transposon family encodes diverse programmable RNA-guided endonucleases. Science 374, 57–65 (2021).

4. T. Karvelis, et al., Transposon-associated TnpB is a programmable RNA-guided DNA endonuclease. Nature 599, 692–696 (2021).

5. M. Saito, et al., Fanzor is a eukaryotic programmable RNA-guided endonuclease. Nature (2023) 10.1038/s41586-023-06356-2.

6. J. A. Doudna, The promise and challenge of therapeutic genome editing. Nature 578, 229–236 (2020).

7. A. Pickar-Oliver, C. A. Gersbach, The next generation of CRISPR-Cas technologies and applications. Nat. Rev. Mol. Cell Biol. 20, 490–507 (2019).

8. F. J. M. Mojica, C. Díez-Villaseñor, J. García-Martínez, C. Almendros, Short motif sequences determine the targets of the prokaryotic CRISPR defence system. Microbiology 155, 733–740 (2009).

9. M. Ciciani, et al., Automated identification of sequence-tailored Cas9 proteins using massive metagenomic data. Nat. Commun. 13, 6474 (2022).

10. J. R. Davis, et al., Efficient in vivo base editing via single adeno-associated viruses with size-optimized genomes encoding compact adenine base editors. Nat Biomed Eng 6, 1272–1283 (2022).

11. D. Wang, F. Zhang, G. Gao, CRISPR-Based Therapeutic Genome Editing: Strategies and In Vivo Delivery by AAV Vectors. Cell 181, 136–150 (2020).

12. G. Gasiunas, et al., A catalogue of biochemically diverse CRISPR-Cas9 orthologs. Nat. Commun. 11, 5512 (2020).

13. K. S. Makarova, et al., Evolutionary classification of CRISPR-Cas systems: a burst of class 2 and derived variants. Nat. Rev. Microbiol. 18, 67–83 (2020).

14. B. Al-Shayeb, et al., Diverse virus-encoded CRISPR-Cas systems include streamlined genome editors. Cell 185, 4574–4586.e16 (2022).

15. R. Nakagawa, et al., Engineered Campylobacter jejuni Cas9 variant with enhanced activity and broader targeting range. Commun Biol 5, 211 (2022).

16. L. Zhang, et al., Author Correction: AsCas12a ultra nuclease facilitates the rapid generation of therapeutic cell medicines. Nat. Commun. 12, 4500 (2021).

17. D. Y. Kim, et al., Efficient CRISPR editing with a hypercompact Cas12f1 and engineered guide RNAs delivered by adeno-associated virus. Nat. Biotechnol. 40, 94–102 (2022).

18. F. A. Ran, et al., In vivo genome editing using Staphylococcus aureus Cas9. Nature 520, 186–191 (2015).

19. A. Casini, et al., A highly specific SpCas9 variant is identified by in vivo screening in yeast. Nat. Biotechnol. 36, 265–271 (2018).

20. E. Kim, et al., In vivo genome editing with a small Cas9 orthologue derived from Campylobacter jejuni. Nat. Commun. 8, 14500 (2017).

21. F. Li, et al., Comparison of CRISPR/Cas Endonucleases for in vivo Retinal Gene Editing. Front. Cell. Neurosci. 14, 570917 (2020).

22. J. L. Schmid-Burgk, et al., Highly Parallel Profiling of Cas9 Variant Specificity. Mol. Cell 78, 794–800.e8 (2020).

23. S. Q. Tsai, et al., GUIDE-seq enables genome-wide profiling of off-target cleavage by CRISPR-Cas nucleases. Nat. Biotechnol. 33, 187–197 (2015).

24. R. M. Daer, J. P. Cutts, D. A. Brafman, K. A. Haynes, The Impact of Chromatin Dynamics on Cas9-Mediated Genome Editing in Human Cells. ACS Synth. Biol. 6, 428– 438 (2017).

25. X. Wu, et al., Genome-wide binding of the CRISPR endonuclease Cas9 in mammalian cells. Nat. Biotechnol. 32, 670–676 (2014).

26. X. Chen, et al., Probing the impact of chromatin conformation on genome editing tools. Nucleic Acids Res. 44, 6482–6492 (2016).

27. R. M. Yarrington, S. Verma, S. Schwartz, J. K. Trautman, D. Carroll, Nucleosomes inhibit target cleavage by CRISPR-Cas9 in vivo. Proc. Natl. Acad. Sci. U. S. A. 115, 9351–9358 (2018).

28. R. S. Isaac, et al., Nucleosome breathing and remodeling constrain CRISPR-Cas9 function. Elife 5 (2016).

29. M. A. Horlbeck, et al., Nucleosomes impede Cas9 access to DNA in vivo and in vitro. Elife 5 (2016).

30. I. Strohkendl, et al., Inhibition of CRISPR-Cas12a DNA targeting by nucleosomes and chromatin. Sci Adv 7 (2021).

31. X. Ding, et al., Improving CRISPR-Cas9 Genome Editing Efficiency by Fusion with Chromatin-Modulating Peptides. CRISPR J 2, 51–63 (2019).

32. L. Dong, et al., An anti-CRISPR protein disables type V Cas12a by acetylation. Nat. Struct. Mol. Biol. 26, 308–314 (2019).

33. Y. Niu, et al., A Type I-F Anti-CRISPR Protein Inhibits the CRISPR-Cas Surveillance Complex by ADP-Ribosylation. Mol. Cell 80, 512–524.e5 (2020).

34. X. Kang, et al., Reversible regulation of Cas12a activities by AcrVA5-mediated acetylation and CobB-mediated deacetylation. Cell Discov 8, 45 (2022).

35. T. Ergünay, et al., Sumoylation of Cas9 at lysine 848 regulates protein stability and DNA binding. Life Sci Alliance 5 (2022).

36. B. P. Kleinstiver, et al., Engineered CRISPR-Cas9 nucleases with altered PAM specificities. Nature 523, 481–485 (2015).

37. B. P. Kleinstiver, et al., Engineered CRISPR-Cas12a variants with increased activities and improved targeting ranges for gene, epigenetic and base editing. Nat. Biotechnol. 37, 276–282 (2019).

38. I. M. Slaymaker, et al., Rationally engineered Cas9 nucleases with improved specificity. Science 351, 84–88 (2016).

39. M. Yamada, et al., Crystal Structure of the Minimal Cas9 from Campylobacter jejuni Reveals the Molecular Diversity in the CRISPR-Cas9 Systems. Mol. Cell 65, 1109– 1121.e3 (2017).

40. S. Hirano, et al., Structural basis for the promiscuous PAM recognition by Corynebacterium diphtheriae Cas9. Nat. Commun. 10, 1968 (2019).

41. A. Das, et al., The molecular basis for recognition of 5’-NNNCC-3’ PAM and its methylation state by Acidothermus cellulolyticus Cas9. Nat. Commun. 11, 6346 (2020).

42. P. Pausch, et al., DNA interference states of the hypercompact CRISPR-CasΦ effector. Nat. Struct. Mol. Biol. 28, 652–661 (2021).

43. G. Xiang, et al., Evolutionary mining and functional characterization of TnpB nucleases identify efficient miniature genome editors. Nat. Biotechnol. (2023) 10.1038/s41587-023-01857-x.

44. F. Storici, M. A. Resnick, The delitto perfetto approach to in vivo site-directed mutagenesis and chromosome rearrangements with synthetic oligonucleotides in yeast. Methods Enzymol. 409, 329–345 (2006).

45. R. D. Gietz, R. H. Schiestl, High-efficiency yeast transformation using the LiAc/SS carrier DNA/PEG method. Nat. Protoc. 2, 31–34 (2007).

46. M. Martin, Cutadapt removes adapter sequences from high-throughput sequencing reads. EMBnet J. 17, 10 (2011).

47. B. Langmead, S. L. Salzberg, Fast gapped-read alignment with Bowtie 2. Nat. Methods 9, 357–359 (2012).

48. S. Valentini, T. Fedrizzi, F. Demichelis, A. Romanel, PaCBAM: fast and scalable processing of whole exome and targeted sequencing data. BMC Genomics 20, 1018 (2019).

49. E. K. Brinkman, T. Chen, M. Amendola, B. van Steensel, Easy quantitative assessment of genome editing by sequence trace decomposition. Nucleic Acids Res. 42, e168 (2014).

50. G. Petris, et al., Hit and go CAS9 delivered through a lentiviral based self-limiting circuit. Nat. Commun. 8, 15334 (2017).

51. C. L. Nobles, et al., iGUIDE: an improved pipeline for analyzing CRISPR cleavage specificity. Genome Biol. 20, 14 (2019).

52. J. Jumper, et al., Highly accurate protein structure prediction with AlphaFold. Nature 596, 583–589 (2021).

53. M. Varadi, et al., AlphaFold Protein Structure Database: massively expanding the structural coverage of protein-sequence space with high-accuracy models. Nucleic Acids Res. 50, D439–D444 (2022).

54. L. Holm, Dali server: structural unification of protein families. Nucleic Acids Res. 50, W210–W215 (2022).

55. E. F. Pettersen, et al., UCSF ChimeraX: Structure visualization for researchers, educators, and developers. Protein Sci. 30, 70–82 (2021).

