## Supplementary information for "Eukaryotic-driven directed evolution of Cas9 nucleases"

**Supplementary Figure 1**

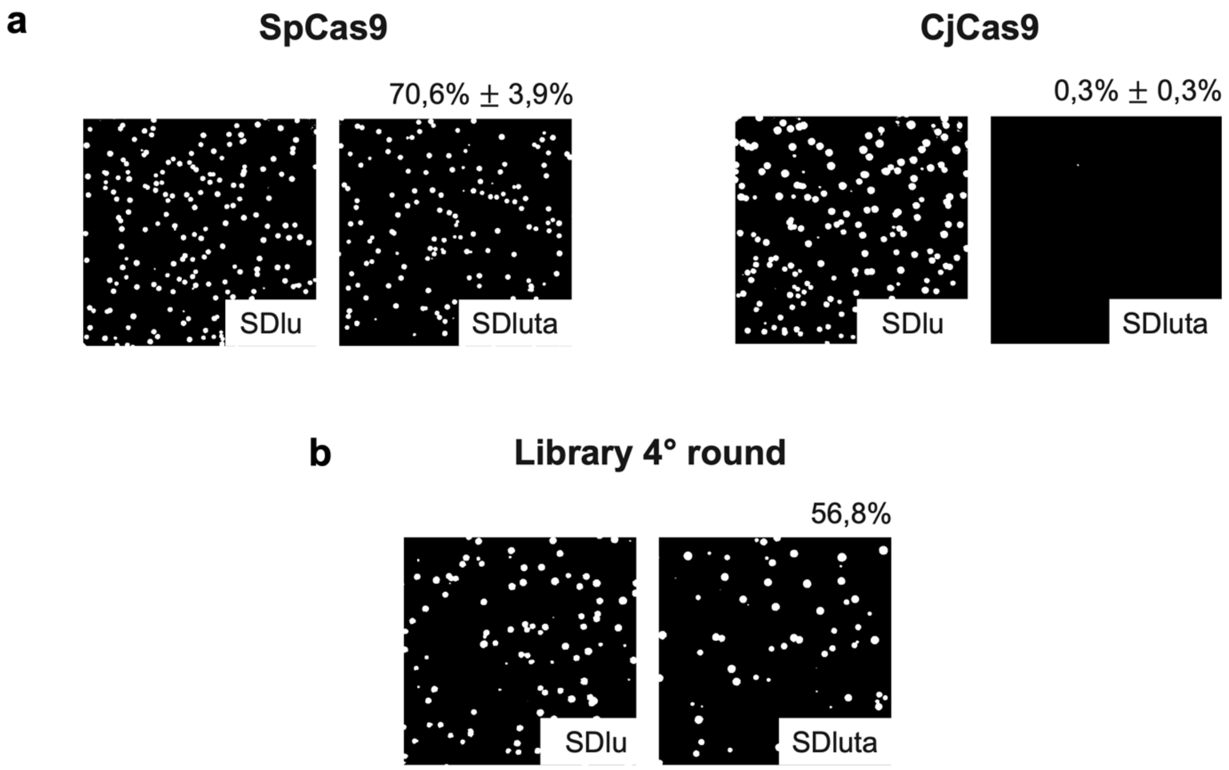


**Supplementary Figure 1. Yeast cultures from the auxotrophic evolution platform**. (**a**) Representative plates obtained by transforming yeasts with SpCas9 and CjCas9 plasmids. Reported percentages are means ± standard deviation of colony survival for n=3 biologically independent replicates. Survival rates are determined as ratios of colonies growing in SDluta (cleaved cassettes) and SDlu (transformants). (**b**) Representative plates showing the survival rate of yeast colonies after transformation with the 4th round library as a PCR amplicon. The mean percentage of survival is reported above the panels.

### Supplementary Figure 2

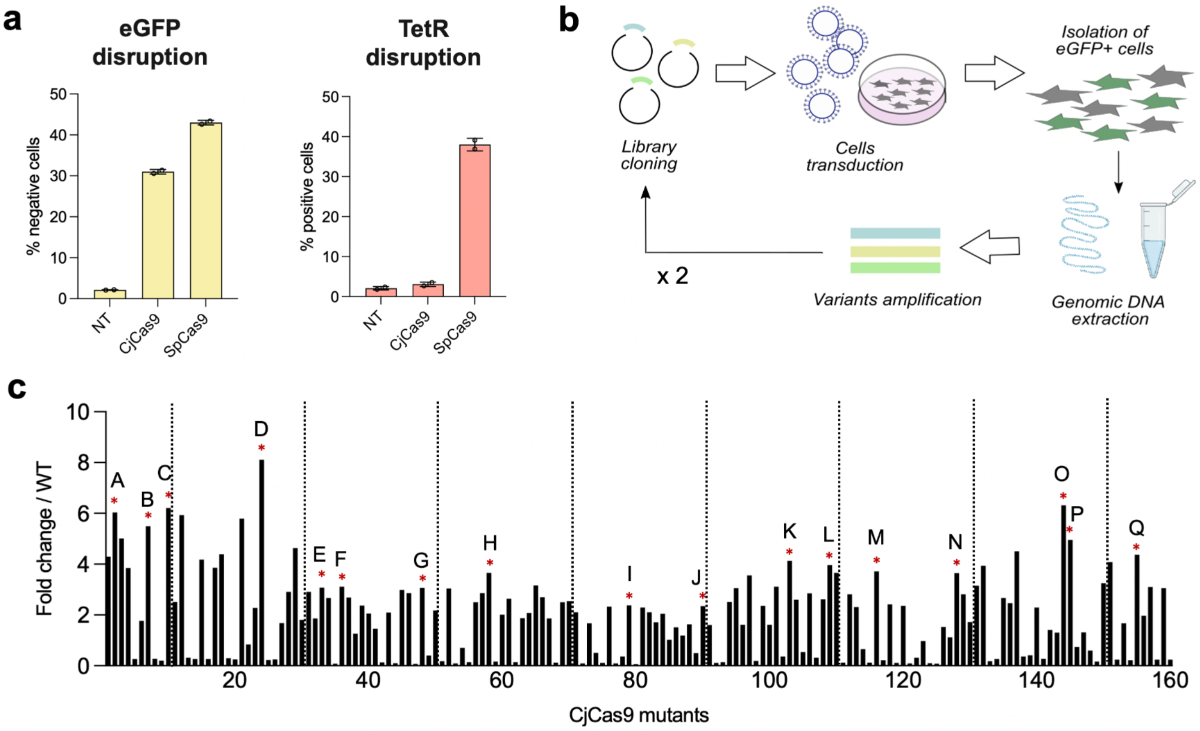


**Supplementary Figure 2. TetR-EGFP reporter screening platform.** (**a**) Validation of the TetR-EGFP reporter cell line with CjCas9 and SpCas9, compared with EGFP disruption assay. Data shown as percentages of EGFP negative or positive cells obtained by FACS analysis 6 days after transfection with CjCas9 and SpCas9 targeting EGFP or TetR. Data are reported as mean ± standard deviation of n=2 biologically independent samples. Individual values are represented as empty circles. (**b**) Schematic representation of the screening rounds in the TetR-EGFP reporter cell line. Variants are amplified at each round and cloned in lentiviral vectors for transduction of the reporter cell line and isolation of EGFP positive cells by sorting. The process was repeated twice. (**c**) Test of cleavage activity of n=160 randomly picked CjCas9 mutants from the 4th round library after enrichment screening in mammalian cells using the TetR reporter system. Data are reported as fold changes of the percentages of EGFP positive cells with respect to CjCas9 WT; values were obtained by FACS analysis 6 days after transfection of the reporter cell line with each variant plasmid. Dotted lines distinguish different groups of transfections. Considering the high variability between different groups, the best performing variants of each group (pointed out by red asterisks and letters) were selected for further characterization.

**Supplementary Figure 3**

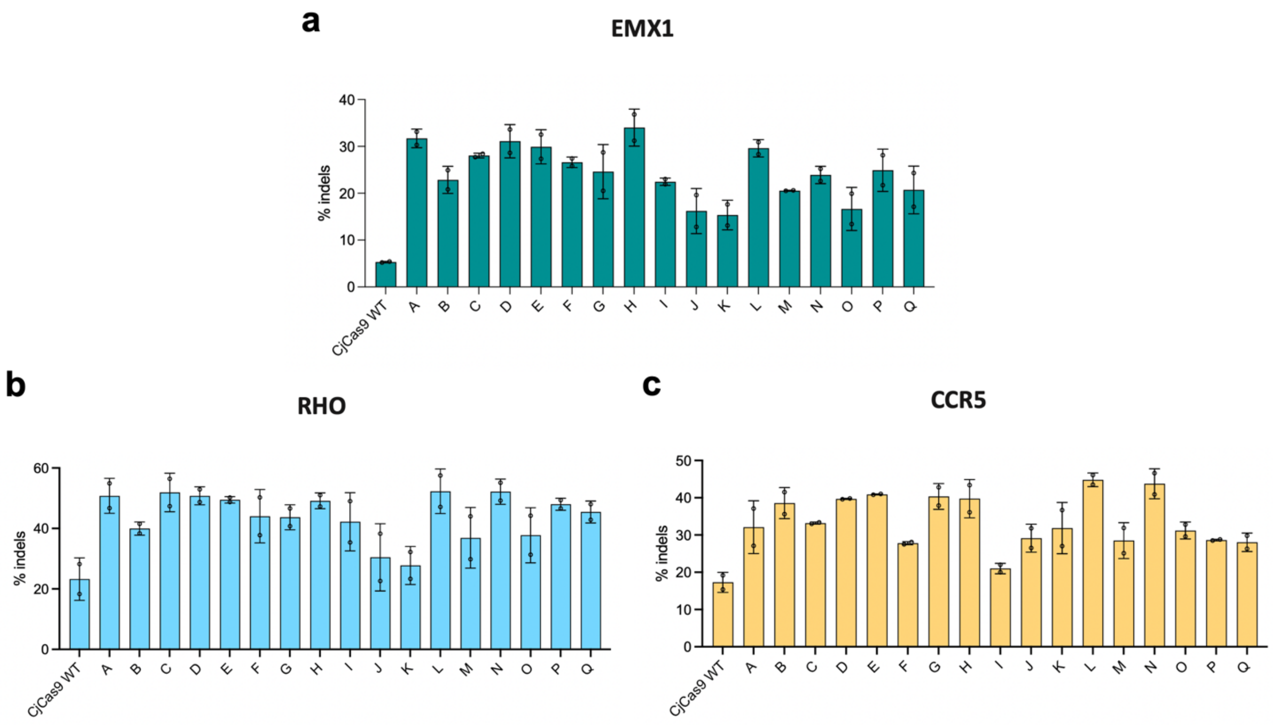


**Supplementary Figure 3. Editing activity of selected variants in endogenous loci of HEK293 cells.** (**a**) DNA cleavage of the *EMX1* endogenous locus with CjCas9 and selected variants (mutants A to Q) after transient transfections. (**b**) DNA cleavage of *RHO* endogenous locus with CjCas9 and selected variants (mutants A to Q). (**c**) DNA cleavage of *CCR5* endogenous locus with CjCas9 and selected variants (mutants A to Q). Data reported as mean ± standard deviation of indels percentages of n=2 biologically independent samples. Individual values are represented as empty circles.

**Supplementary Figure 4**

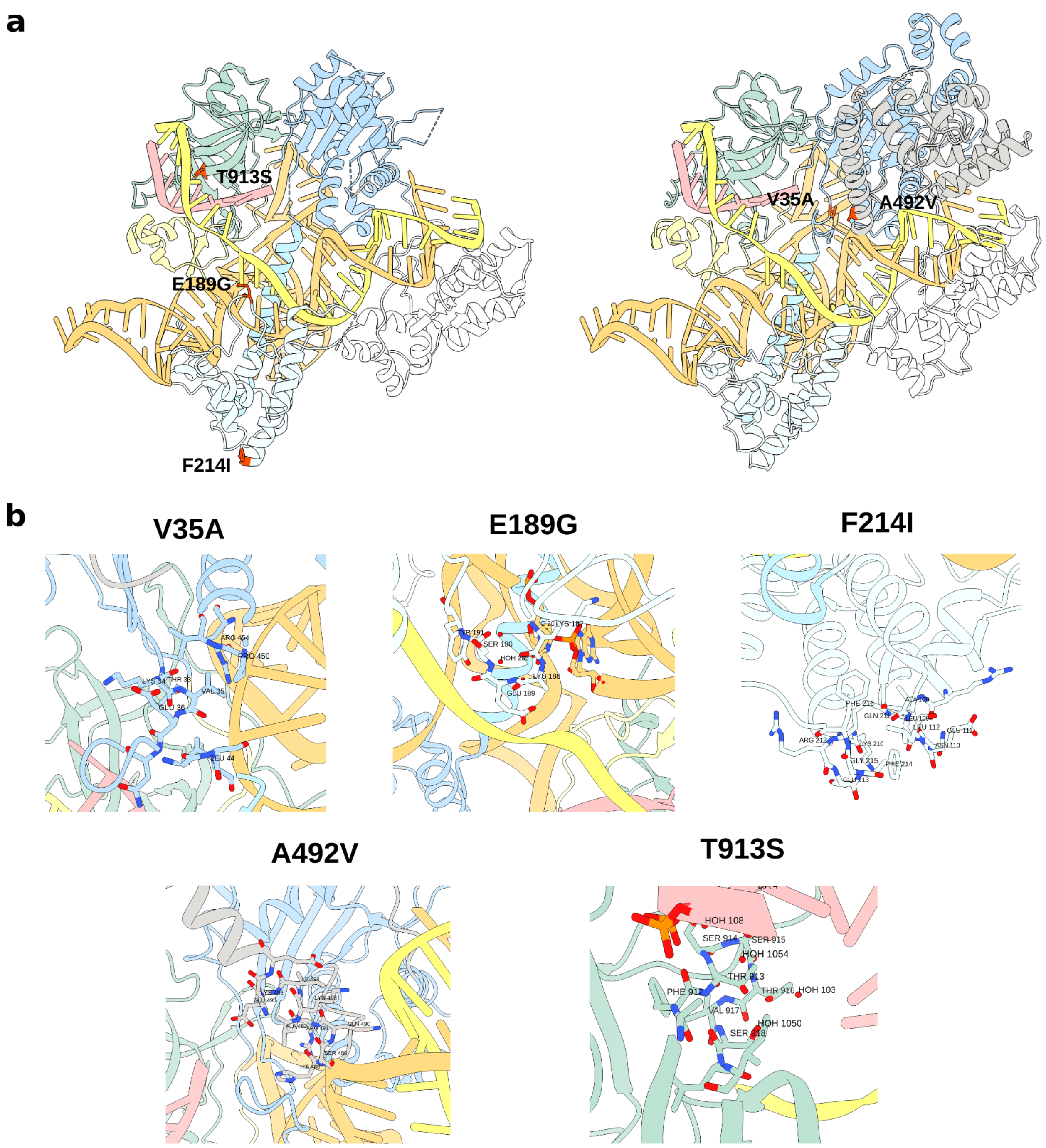


**Supplementary Figure. 4. 3D position of the mutations in UltraCjCas9.** (**a**) Representation of key residues mutated in UltraCjCas9 (mutant H). Amino acids 189, 214 and 913 were highlighted in red in the crystal structure of CjCas9 (PDB ID 5X2H) using

UCSF ChimeraX. Residues 35 and 492, which are missing from the crystal structure, are highlighted in red in the predicted structure of CjCas9 retrieved from the AlphaFold Protein Structure Database (accession Q0P897). CjCas9 domains are reported using the color scheme of the protein structure shown in **Figure 3a**. **b)** Zoomed-in view of each mutated residue and surrounding area.

### Supplementary Figure 5

**a**

| VRYAC | |
|---|---|
| HEK site 2 | AATAC |
| HBB | |
| BCR | CATAC |
| FAS1 | CGTAC |
| HBG1 | GATAC |
| CCR5 | GGTAC |
| TRAC | |
| RHO | AGTAC |
| HPRT1 | |
| EMX1 | AGCAC |
| FANCF | |
| CHR8 | AACAC |
| HEK site 4 | TACAC |
| ZSCAN2 | CGCAC |
| HEK site 1 | GGCAC |
| TRBC1 | |
| IL2RG | GACAC |
| VEGFA2 | |
| PD1 | CACAC |
| VEGFA3 | |

| Not VRYAC | |
|---|---|
| CHR6 | AACAT |
| ATM | AACAA |
| HEK site 3 | AACAG |

**b**

| | |
|---|---|
| **TRBC** | CAGCGCACACCATGAAGGAGAATTGGGCAC |
| **TRAC** | TCACTGGATTTAGAGTCTCTCAGCTGGTAC |
| **HEK site 1** | GGGCAAGTGCTGTCTGCCATGCGCCGGCAC |
| **HEK site 2** | AACTTCCCAAGTGAGAAGCCAGTGGAATAC |
| **FANCF** | CCGAATTCCCCAGAAGCCAGTGGACTAGCAC |

**c**

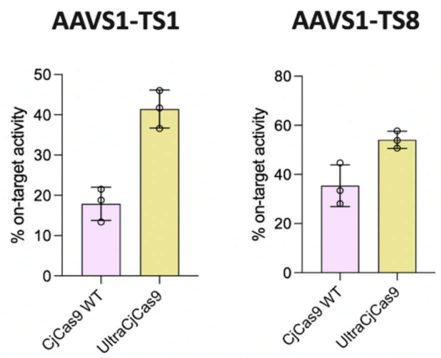


**Supplementary Figure 5**. (**a**) List of PAM sequences selected to design gRNAs targeting genomic loci. NNNVRYAC and not NNNVRYAC PAM sequences were chosen to widely evaluate the editing efficiency of UltraCjCas9. gRNAs are listed in **Supplementary Table 3**. (**b**) DNA sequences representing the relative positions of SpCas9 and UltraCjCas9 gRNAs targeting 5 selected genomic loci. PAM sequences of SpCas9 are underlined in gray, PAM sequences of UltraCjCas9 in yellow. (**c**) DNA on-target cleavage of *AAVS1-TS1* and *AAVS1-TS8* endogenous loci with CjCas9 WT and UltraCjCas9 after transfection. Data reported as mean ± standard deviation of indels percentages of n=3 biologically independent samples. Individual values are represented as empty circles.
